# Sex-specific effect of antenatal Zika virus infection on murine fetal growth, placental nutrient transporters, and nutrient sensor signaling pathways

**DOI:** 10.1101/2023.03.30.534997

**Authors:** Daniela Pereira-Carvalho, Alessandra Cristina Chagas Valim, Cherley Borba Vieira Andrade, Enrrico Bloise, Ariane Fontes Dias, Veronica Muller Oliveira Nascimento, Rakel Kelly Silva Alves, Felipe Lopes Brum, Inácio Gomes Medeiros, Sharton Vinicius Antunes Coelho, Luciana Barros Arruda, Adriane Regina Todeschini, Wagner Barbosa Dias, Tania Maria Ortiga-Carvalho

## Abstract

Maternal Zika virus (ZIKV) infection during pregnancy can associate with severe intrauterine growth restriction (IUGR), placental damage, and metabolism disturbance, as well as newborn neurological abnormalities. Here, we investigated whether maternal ZIKV infection affects placental nutrient transporters and nutrient-sensitive pathways. Immunocompetent (C57BL/6) mice were injected with Low (10^3^ PFU-ZIKV_PE243_) and High (5×10^7^ PFU-ZIKV_PE243_) ZIKV titers at gestational day (GD) 12.5, for tissue collection at GD18.5 (term). Feto-placental growth of male fetuses was dramatically affected by ZIKV, whereas no differences were observed in female fetuses. ZIKV promoted increased expression of glucose transporter type 1 (*Slc2a1*/Glut1) and decreased levels of glucose-6-phosphate in female placentas, with no differences in amino-acid transport potential. In contrast, glucose transport in male placentas was not affected by ZIKV, whilst a decreased placental protein expression of sodium-coupled neutral amino acid 2 (Snat2) was detected in the male low-dose ZIKV-infected group. There were also sex-dependent differences in the hexosamine biosynthesis pathway (HBP) and O-GlcNAcylation in ZIKV infected pregnancies, showing that ZIKV can cause disturbance in the nutrient handling in the placental tissue. Our findings thus identify relevant molecular alterations in the placenta caused by maternal ZIKV infection related to nutrient transport and availability. Notably, our results suggest that female and male placentas adopt different strategies to cope with the altered metabolic state caused by ZIKV. This may have relevance for understanding the effects of congenital Zika syndrome and could potentially assist future therapeutic strategies.

**Author Summary:** The Zika virus (ZIKV) has emerged as a major global health concern in the past decade. ZIKV infection during pregnancy can cause infants to be born with microcephaly and fetal growth restriction, among other pregnancy complications. Currently, the number of cases of ZIKV disease declined onwards globally. However, transmission persists at low levels in several countries in the Americas and other endemic regions, with neither a licensed vaccine nor an antiviral drug available for prevention and treatment. Here, we use a mice model of maternal ZIKV infection to analyze placental nutrient transporters and nutrient-sensitive pathways as a potential link to the complications related to congenital ZIKV infection. We found that feto-placental growth of male fetuses was dramatically affected by ZIKV, whereas no differences were observed in female fetuses. We also found that placental nutrient transporters and nutrient-sensitive pathways were altered in response to ZIKV infection, depending on the fetal sex. Our study presents relevant molecular alterations caused by maternal ZIKV infection and suggests that female and male placentas adopt different strategies in response to the altered environment caused by ZIKV. Our observations may have relevance for understanding the effects of ZIKV infection and could potentially assist future therapeutic strategies.

## Introduction

Zika virus (ZIKV) is an Aedes-borne arbovirus belonging to the *Flaviviridae* family of positive, single-stranded, enveloped RNA viruses that has emerged as a major global health concern in the past decade. ZIKV infection in pregnant women has been linked to mild to severe neonatal abnormalities, such as microcephaly and intracranial calcifications, as well as pregnancy loss, intrauterine growth restriction (IUGR), and preterm labor, among other complications [1]. The ZIKV pandemic is currently thought to be “controlled”; however, more than 60% of the world’s population live in regions with suitable environmental conditions for ZIKV spread, with neither a licensed vaccine nor an antiviral drug [2] available for prevention and treatment of the disease.

The mechanisms underlying antenatal ZIKV commitment of fetal growth and developmental trajectories have only been partially elucidated. Similar to other flaviviruses, ZIKV requires a variety of intricate interactions with host factors for successful infection, such as reprogramming of cellular metabolism underpinning ZIKV pathogenesis [3–5]. In the placenta, ZIKV is able to cause lipid reprogramming, mitochondrial dysfunction, and inflammatory immune imbalance, contributing to placental damage [3,6,7]. At the functional level, placental damage can impair essential placental barrier mechanisms that support fetal growth and development, such as unidirectional glucose and amino acid transfer from mother to conceptus [8].

Glucose (GLUT) and sodium-coupled neutral amino acid (SNAT) transporters are key placental nutrient transporters - highly regulated by environmental factors such as oxygen, nutrients and hormones [9,10]. Alterations in extracellular glucose and insulin-like growth factor I (IGF-I) levels can cause changes in the activity and expression of glucose and amino acid transporters [11]. When deregulated, GLUT and SNAT transporters have been linked to pregnancy complications, such as IUGR, and the development of neurological and cardiometabolic diseases later in life [12–14]. In the context of infection, *plasmodium falciparum* (the etiological agent of placental malaria) infestation has been shown to decrease GLUT1 in the term human placenta [15]. Additionally, ZIKV infection was linked with an increase in glucose uptake and GLUT3 expression human in first-trimester cytotrophoblast cells [16]. Demonstrating that placental nutrient uptake and metabolism is impacted by common infective agents.

In this connection, the hexosamine biosynthesis pathway (HBP) is a neglected anabolic branch of glucose metabolism that uses glucose and glutamine to generate UDP-GlcNAc as the final product. Glutamine-fructose amino transferase (GFAT) is HBP’s rate-limiting enzyme, whereas UDP-GlcNAc is considered a central metabolic sensor since it requires inputs from carbohydrate, amino acid, nucleotide and fatty acid metabolism. UDP-GlcNAc is the donor substrate of O-GlcNAc transferase (OGT), the enzyme responsible for the addition of O-GlcNAc to serine/threonine residues of target intracellular proteins, while O-GlcNAcase (OGA) is the enzyme responsible for its removal [17]. O-GlcNAcylation is a highly dynamic and ubiquitous posttranslational modification that modulates protein function in a nutrient-sensitive way, linking cell signaling to metabolism [17]. Cellular UDP-GlcNAc levels and global O-GlcNAcylation are coordinated and are highly responsive to glucose availability, making O-GlcNAcylation well suited nutrient-sensor [18]. HBP and O-GlcNAcylation are central in controlling immune cell function, consequently having key regulatory effects on immune responses against infections [19]. The molecular mechanisms involving O-GlcNAcylation were investigated using different virus infection models, including HPV [20,21], HBV [22], influenza A virus [23], and HSV [24]. Although these studies demonstrated that viruses are able to modulate O-GlcNAcylation, impacting on autophagy mechanisms and viral replication, the effects of the Zika virus remain unexplored.

In addition, OGT and OGA are highly expressed in the placenta, and key placental proteins are OGT targets. Placental OGT expression has been linked to long-term metabolic and neurodevelopmental programming [25–28]. For instance, studies in mice showed that aberrant O-GlcNAcylation causes reduced placental vasculature development, insulin tolerance, and changes in body weight [27–29]. An OGT knockout mouse model showed important changes in the expression patterns of hypothalamic genes during intrauterine development [25,26].

A previous study by our group showed that maternal exposure to ZIKV affects placental function, including placental ultrastructure and ABC transporter protein expression, even in the absence of vertical transmission and that these effects are dependent on viral infective titers and maternal immune status [7]. Despite their importance to fetal growth and development and their role in response to infections, there is limited information about GLUT and SNAT transporter expression profiles and OGT nutrient-sensing pathway activity in the placenta of ZIKV-infected dams. Therefore, in the present study, we hypothesize that maternal exposure to ZIKV affects fetal-placental growth, the expression of placental nutrient transporters and nutrient distribution, as well as placental O-GlcNAcylation, and specific components of the HBP, in a sex-specific manner.

## Results

### Antenatal ZIKV infection alters fetal-placental growth in a sex-specific manner

Pregnant mice were inoculated at GD12.5 with a Low Dose (LD, 10^3^ PFU) or High Dose (HD, 5×10^7^ PFU) of ZIKVPE243 or with conditioned medium from noninfected C6/36 cells as controls (Fig 1A). To investigate whether maternal ZIKV infection affects placental efficiency and fetal biometry at term, fetuses and placentas were weighed at GD 18.5 (considered term in C57BL/6). No significant difference was observed in the weight of female fetuses and their corresponding placentas, as well as in their placental efficiency (Fig 1B). The fetal head weight of females in the LD group was significantly reduced (-16.02%, p=0.041) compared to corresponding control group (Fig 1B). In contrast, fetal-placental growth was dramatically reduced in male fetuses. The fetal weight was significantly reduced in both infected groups (LD: -13.8%, p=0.002; HD: -20.5%, p=0.032) compared to control male fetuses (Fig 1D). Additionally, placentas from the infected groups exhibited reduced weight (LD: -20,92% p=0.033, HD: -15,21%, p=0.001) compared to the correspondent control group, with no significant difference in placental efficiency (Fig 1D). We therefore investigated whether maternal ZIKV infection would cause placental remodeling, affecting the two main functionally distinguishable regions of the rodent placenta. There were no differences in the areas of the Lz and Jz or in the total placental area when the ZIKV-infected groups were compared to the control group in either female (Fig 1C) or male (Fig 1E) placentas.

**Figure 1.**
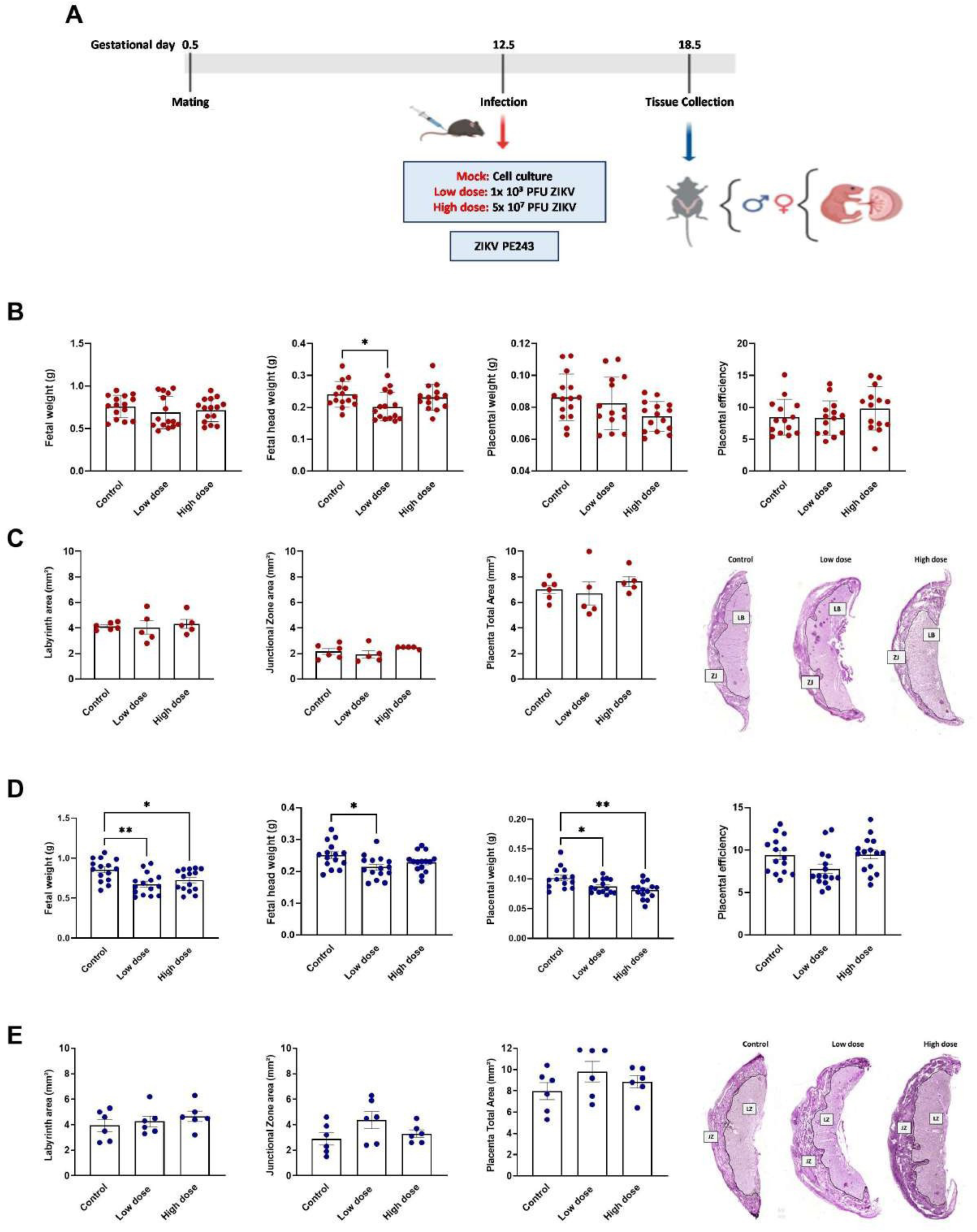
Impact of ZIKV infection on fetal-placental growth. **(A)** Experimental design. **(B)** Fetal weight, fetal head weight, placental weight, and placental efficiency in female fetuses. **(C)** Quantification of the areas of the Lz, Jz, and total placental area in females and representative PAS-stained images showing the delimitation of placental areas. **(D)** Fetal weight, fetal head weight, placental weight, and placental efficiency in male fetuses. **(E)** Quantification of the areas of the Lz, Jz, and total placental area in males and representative PAS-stained images showing the delimitation of placental areas. Data are displayed as the mean + SEM. and analyzed by one-way ANOVA with Tukey post hoc comparisons. (n=15). *p < 0.05, **p < 0.005.

### Maternal ZIKV infection induces changes in placental amino acid and glucose transport

To investigate whether changes in fetal and placental growth may be related to alterations in placental transport of amino acids and glucose, we evaluated the expression of key glucose and amino acid transporters and the levels of glucose-6-phosphate and glutamine in placental tissue. In particular, the mRNA expression of key transporters for glucose (*Slc2a1* and *Slc2a3*) and amino acids (*Slc38a1* and *Slc38a2*) was quantified in the placenta by RT‒ qPCR. Additionally, the protein expression of the transporters Glut1 and Snat2 was quantified by immunohistochemistry in both the placenta Lz and Jz areas. Levels of glucose-6-phosphate and glutamine was analyzed by MALDI-MSI.

In placentas from female fetuses, these analyses showed that the expression of *Slc2a1* mRNA in the HD group was 62% greater (p=0.047) than in the control group, while this difference was not observed for the LD group (Fig 2A). No differences were observed in *Slc2a3* mRNA expression between the experimental groups (Fig 2A). In addition, no differences were found in *Slc38a1* and *Slc38a2* mRNA expression in female placentas (Fig 2B). At the protein expression level, the expression of Glut1 in the Lz of female placentas was greater in the ZIKV-infected groups (LD: +73.40%, p=0.007; HD: +73.38%, p=0.007) than in the control group (Fig 2C). However, Glut1 expression did not vary among groups in the Jz region of female placentas (Supplementary Fig 1A). We evaluated the protein expression of Snat2 in female placental tissue and found no differences between the Zika-infected and control groups in either the Lz (Fig 2D) or Jz (Supplementary Fig 1B) regions. Interestingly, levels of glucose-6-phosphate was reduced in the ZIKV-infected groups compared to the control placentas (Fig 2E), whereas the levels of glutamine in the female placenta was similar between groups (Fig 2E).

**Figure 2.**
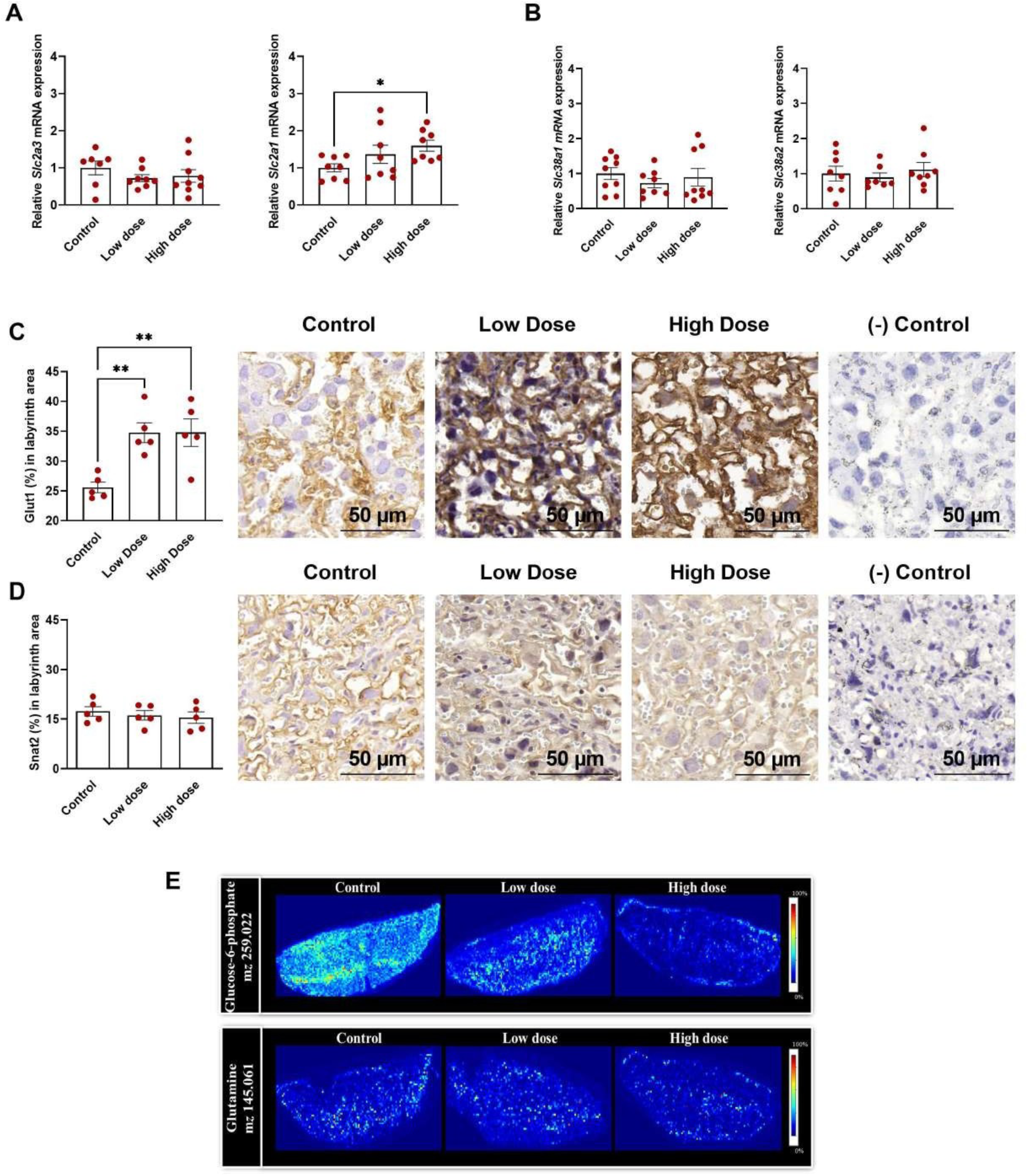
Nutrient transport in control and ZIKV-infected female placentas. **(A)** Slc2a1 and Slc2a3 mRNA expression by qPCR (n=8-10). **(B)** Slc38a1 and Slc38a2 mRNA expression by qPCR (n=8-10). (**C)** Quantification and representative photomicrographs of Glut1 expression in the placental transport zone by immunohistochemistry (n=5). **(D)** Quantification and representative photomicrographs of Snat2 expression in the placental transport zone by immunohistochemistry (n=5). **(E**) Matrix-assisted laser desorption/ionization mass spectrometry imaging analysis showing the distribution of glucose-6-phosphate by 259.022 m/z localization in placental tissue and the distribution of glutamine by 145.061 m/z localization in placental tissue (n=5). Data are displayed as the mean + SEM. and analyzed by one-way ANOVA with Tukey post hoc comparisons. *p < 0.05, **p < 0.005.

The same parameters were evaluated in placentas from males, but the results showed different responses. The mRNA expression of key glucose transporters, *Slc2a1* and *Slc2a3*, was not significantly different between the experimental groups (Fig 3A). Additionally, the mRNA expression of the amino acid transporters *Slc38a1* and *Slc38a2* was not significantly different between the ZIKV-infected and control groups (Fig 3B). In addition, the protein expression of Glut1 in male placentas was not different between the experimental groups in either Lz (Fig 3C) or Jz (Supplementary Fig 1C). While Snat2 protein expression in the Lz of male placentas was lower in the LD group than in the control group (-63.66%, p=0.025), no differences were observed in the comparison with the HD group (Fig 3D). We found no differences between the experimental groups when analyzing Snat2 protein expression in the Jz of male placentas (Supplementary Fig 1D). Meanwhile, the presence of essential molecules in male placentas, glucose-6-phosphate and glutamine placenta were similar between the ZIKV-infected and control groups (Fig 3E).

**Figure 3.**
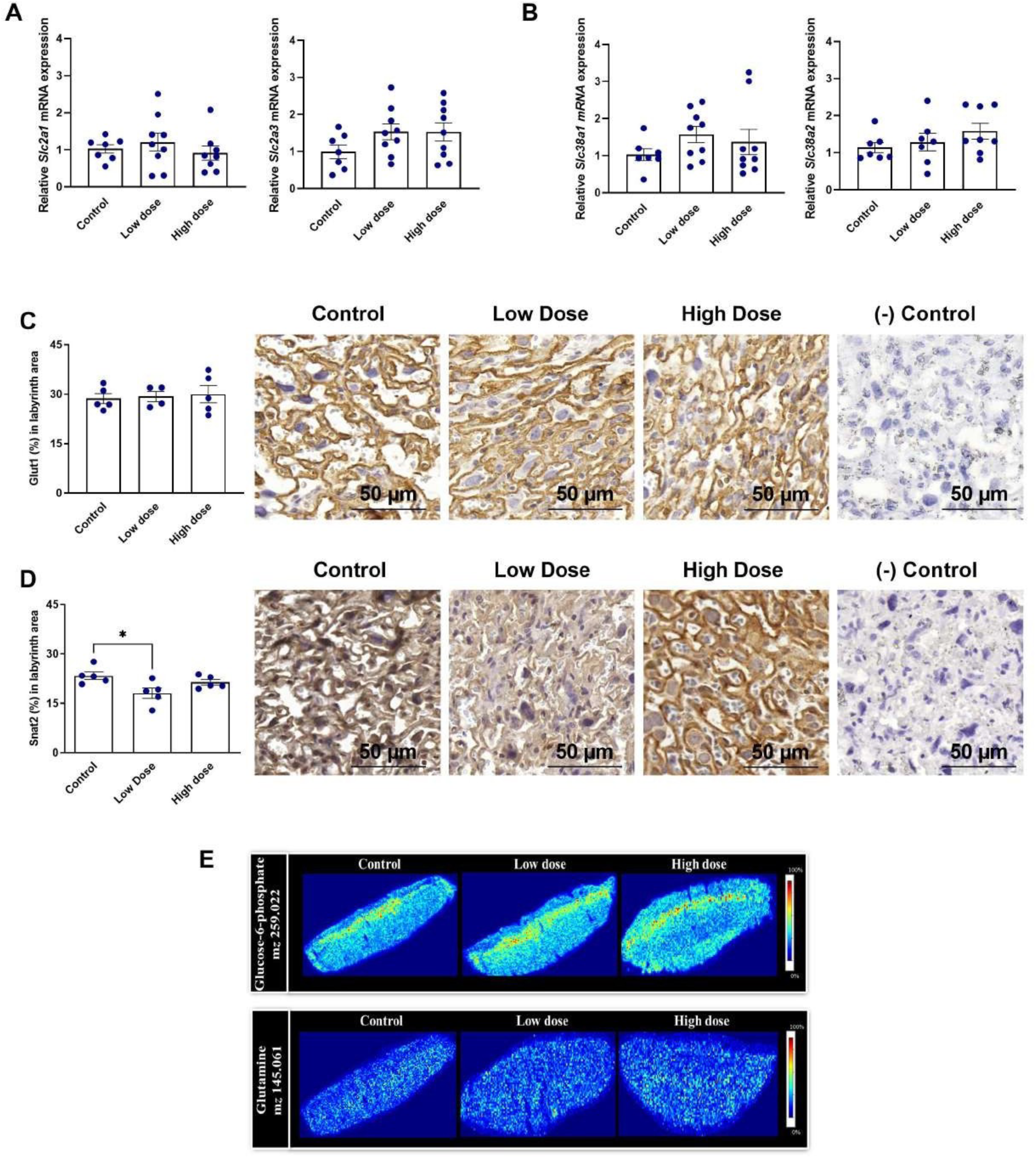
Nutrient transport in control and ZIKV-infected male placentas. **(A)** *Slc2a1* and *Slc2a3* mRNA expression by qPCR (n=8-10). **(B)** *Slc38a1* and *Slc38a2* mRNA expression by qPCR (n=8-10). (**C)** Quantification and representative photomicrographs of Glut1 expression in the placental transport zone by immunohistochemistry (n=5). **(D)** Quantification and representative photomicrographs of Snat2 expression in the placental transport zone by immunohistochemistry (n=5). **(E**) Matrix-assisted laser desorption/ionization mass spectrometry imaging analysis showing the distribution of glucose-6-phosphate by 259.022 m/z localization in placental tissue and the distribution of glutamine by 145.061 m/z localization in placental tissue (n=5). Data are displayed as the mean + SEM. and analyzed by one-way ANOVA with Tukey post hoc comparisons. *p < 0.05.

### Maternal ZIKV infection alters OGT expression and O-GlcNAcylation in female placentas

To gain further information about the changes in fetal growth and nutrient transport and distribution, we evaluated pathways sensitive to nutrient availability. For this, we analyzed the mRNA and protein expression of components of the HBP and O-GlcNAcylation. We also analyzed the presence of UDP-GlcNAc, the final product of the HBP.

The analysis of *Gfpt1* mRNA expression showed no significant difference in female placentas (Fig 4A). However, we found that the protein expression of Gfat1 was lower in the HD (-33.73%, p=0.038) group than in the control group, with no significant changes in the LD group (Fig 4B). Interestingly, analysis of the presence of UDP-GlcNAc showed that there was a lower presence of this product in both ZIKV-infected groups in female placentas (Fig 4D). In addition, *Ogt* mRNA expression was lower in the HD group than in the control group (-42.17%, p=0.038), but the comparison with the LD group did not present significant changes (Fig 4D). Additionally, no differences were observed in *Oga* mRNA expression between the experimental groups (Fig 4D). While the protein expression of OGT and OGA was not different between the groups, the O-GlcNAcylation levels were significantly lower in the LD (-35.12%, p=0.045) and HD (-38.09%, p=0.029) ZIKV-infected groups than in the control group in female placentas (Fig 4E).

**Figure 4.**
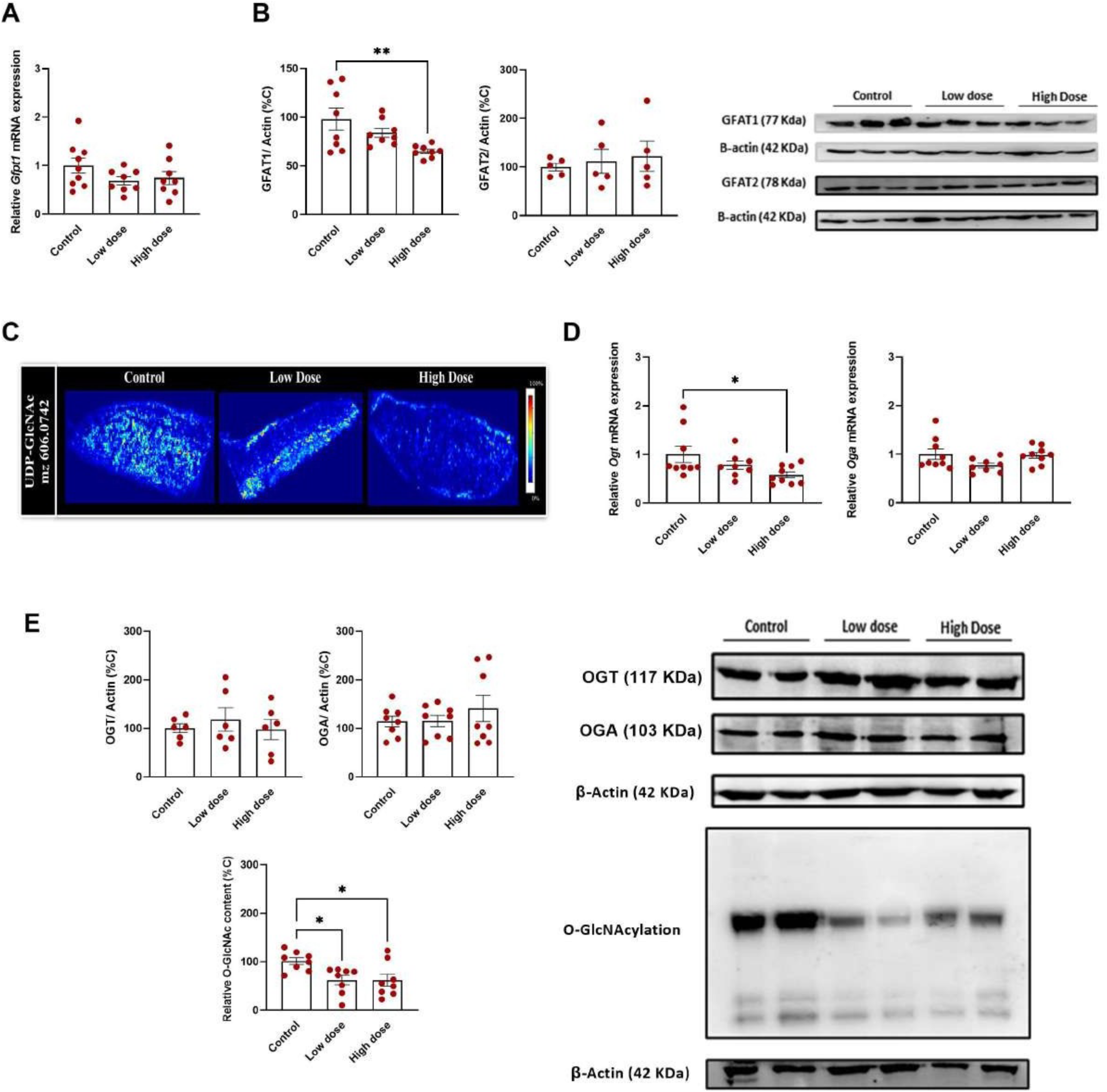
HBP and O-GlcNAcylation in control and ZIKV-infected female placentas. **(A)** *Gfpt1* mRNA expression by qPCR (n=8-10). **(B)** Relative abundance of GFAT1 and GFAT2 (n=5-8). (**C)** Matrix-assisted laser desorption/ionization mass spectrometry imaging analysis showing the distribution of UDP-GlcNAc by 606.0742 m/z localization in placental tissue (n=5). **(D)** *Ogt* and *Oga* mRNA expression by qPCR (n=8-9). **(E**) Relative abundance of OGT, OGA and O-GlcNAcylation (n=6-8). Data are displayed as the mean + SEM. and analyzed by one-way ANOVA with Tukey post hoc comparisons. *p < 0.05, **p < 0.005

In male placentas, the relative *Gfpt1* mRNA expression showed no significant difference between the ZIKV-infected and control groups (Fig 5A). However, Gfat1 protein expression in the HD group was greater than that in the control group (+32.54%, p=0.039), with no significant changes in the LD group (Fig 5B). In addition, the analysis of UDP-GlcNAc in the placenta of males was similar between groups (Fig 5C). Then, we analyzed the mRNA expression of the enzymes related to O-GlcNAcylation. The *Ogt* mRNA HD group was greater than the LD (+49.41%, p=0.038) and control (+48.03%, p=0.031) groups. The *Oga* mRNA expression was also greater in the HD group than the LD (+52.67%, p= 0.030) and control (+16.52%, p=0.048) groups (Fig 5D). The protein expression of OGT and OGA as well as O-GlcNAcylation levels were not different between the experimental groups, showing that O-GlcNAcylation was not affected by ZIKV in males (Fig 5E).

**Figure 5.**
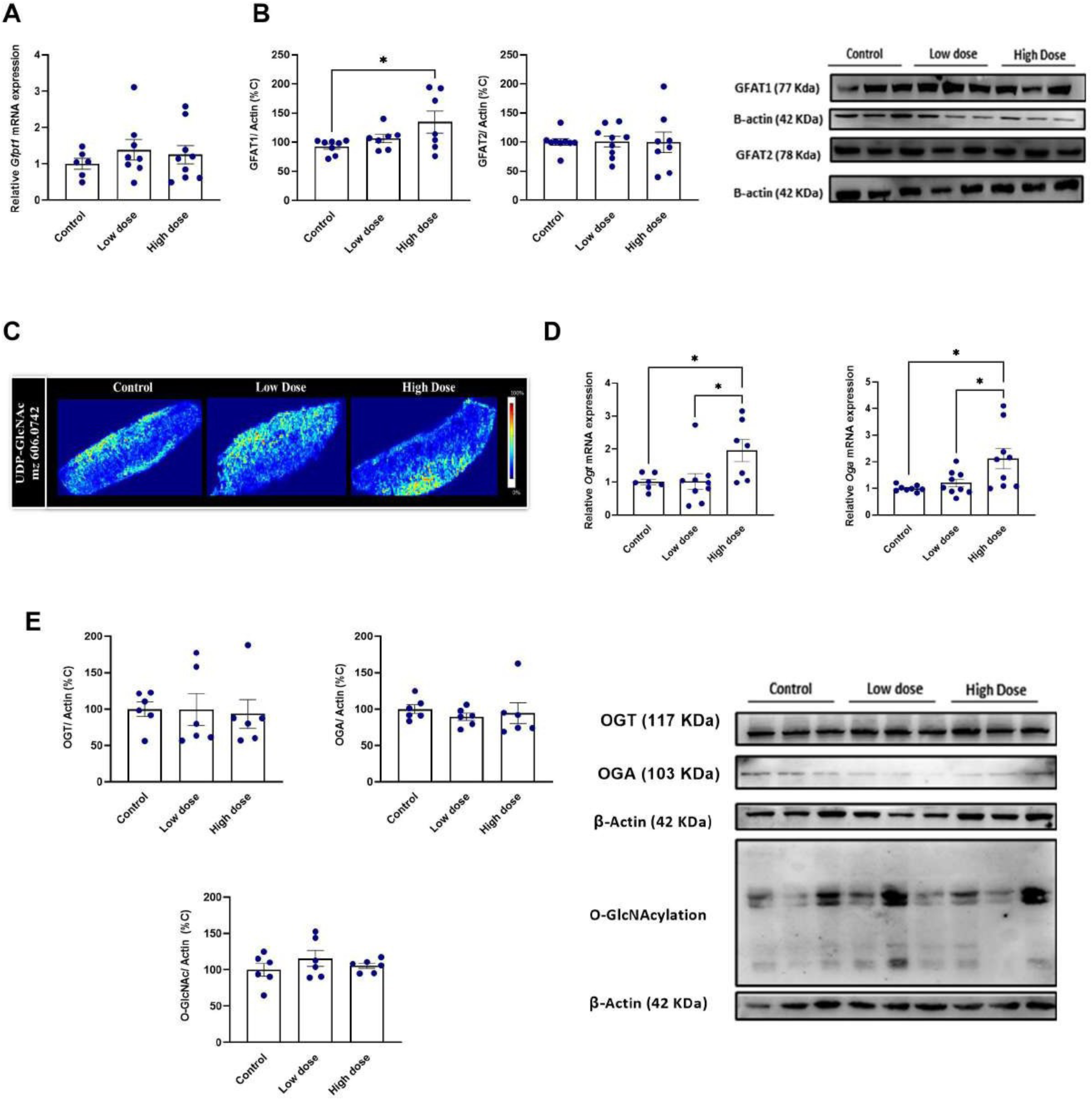
HBP and O-GlcNAcylation in control and ZIKV-infected male placentas. **(A)** *Gfpt1* mRNA expression by qPCR (n=6-8). **(B)** Relative abundance of GFAT1 and GFAT2 (n=7-8). (**C)** Matrix-assisted laser desorption/ionization mass spectrometry imaging analysis showing the distribution of UDP-GlcNAc by 606.0742 m/z localization in placental tissue (n=5). **(D)** *Ogt* and *Oga* mRNA expression by qPCR (n=7-8). **(E**) Relative abundance of OGT, OGA and O-GlcNAcylation (n=6). Data are displayed as the mean + SEM. and analyzed by one-way ANOVA with Tukey post hoc comparisons. *p < 0.05.

## Discussion

In the present study, we investigated the effect of maternal ZIKV infection on placental makers related to nutrient transport and availability using a mouse model of impaired fetal growth and survival with no vertical virus transmission [7]. Herein, we showed that maternal ZIKV infection during pregnancy impacts the litter and their placentas in a sex-specific manner, with variations in nutrient transport and O-GlcNAcylation mechanisms, suggesting that females and males adopt different strategies to cope with the altered placental metabolic state caused by ZIKV infection.

The fetal-placental growth in the female litter was slightly affected by maternal infection. However, fetal-placental growth in males was dramatically affected by ZIKV, independently of viral titers. The reduction in the weight of fetuses and placentas is a known impact of ZIKV infection [30–33]. In fact, studies have demonstrated the potential correlation between fetal and placental weight in mice, as fetal growth relies on a variety of placental functions, such as nutrient transport and hormonal production, that can be affected by the reduction in this organ’s size, impacting fetal growth [34]. It is important to mention that fetal growth restriction during pregnancy enhances the risk for preterm birth, fetal mortality, and the development of chronic diseases later in life through developmental programming [34–36].

Moreover, the capability of ZIKV to induce sex-specific responses has been shown in different models in the literature [37,38]. Interestingly, our study shows that these differences start to be modulated during intrauterine life. The mechanisms by which the placenta supports fetal growth are likely differ between female and male fetuses within the litter, even in normal physiological mouse pregnancies [39]. However, the mechanisms by which these processes are affected by maternal infection have yet to be determined.

Using the present experimental model, our group had previously shown that ZIKV infection impacted proliferation and apoptosis processes in the placental tissue [7]. In addition, in a mouse model of infection with poly (I:C), a synthetic double-stranded RNA viral mimic, it was shown that the infection decreased proliferation in placental Lz, whereas apoptosis in Jz was increased [35,40]. While these studies show that viruses are able to impair placental remodeling, our results did not show differences in placental Lz and Jz areas in either sex.

In our study, the expression of the glucose transporter Glut1 was greater in the exchange (Lz) zone of the infected groups only in female placentas. Glut1 (*Slc2a1*) is the primary placental glucose transporter, as it is the predominant isoform abundantly expressed during early pregnancy and at term in humans and rodents [41,42]. Alterations in the expression of Glut1 (*Slc2a1*) were observed in models with reduced availability of oxygen and nutrients [9,10], as well as in gestational disorders [43–46], and in response to pharmacological treatment [47]. Therefore, placental Glut1 (*Slc2a1*) seems to have great plasticity and plays a role in responses to alterations in the maternal milieu. Interestingly, the presence of glucose-6-phosphate was reduced in the infected groups. There are several positive regulators of Glut expression in the placenta, and low glucose concentration is one of them [48]. In addition, ZIKV infection was linked with an increase in glucose uptake and GLUT3 expression in first-trimester cytotrophoblast cells [16]. These results together may indicate that more glucose is being transported to the female fetuses and are consistent with the maintenance of fetal growth observed in females. Overall, our study shows that the placental increase in Glut1 expression is an adaptation of the female placenta in response to maternal infection in this infection model.

In males, no changes related to glucose transport and glucose-6-phosphate distribution were observed. Nevertheless, the protein expression of Snat2 was lower in the transport zone of the LD group. Low placental Snat2 expression causes fetal growth restriction [12]. Therefore, there is a possible relationship between reduced Snat2 expression and the reduced fetal growth observed in the fetuses of this group. However, further studies must be performed to confirm this hypothesis.

The HBP is a metabolic pathway that generates the “sensing molecule” UDP-GlcNAc. An essential role in the sensing mechanism of the HBP, especially in the context of glucose availability, is assigned to the enzyme catalyzing the first and rate-limiting step of this pathway, fructose-6-phosphate transaminase (GFAT) [49]. In female placentas, the protein expression of GFAT1 was reduced in the HD group. In addition, the presence of UDP-GlcNAc was reduced in both ZIKV-infected groups. Consistent with these findings, the O-GlcNAcylation levels were reduced in the female placentas from the ZIKV-infected groups. Together, these results indicate reduced flow of nutrients into the HBP in female placentas. Additionally, considering the reduced glucose-6-phosphate and UDP-GlcNAc as well as O-GlcNAcylation, there is an indication of nutrient deprivation in the placental tissue. ZIKV infection triggers several adaptations to cellular metabolism [3–5]. Specifically, regarding glucose metabolism, Thaker et al showed that ZIKV infection reprograms glucose metabolism, changing glucose utilization, which is linked to whether cell death occurs [4]. Specifically, in placental tissue, these changes can affect diverse placental functions related to fetal growth and development and interfere with placental intrinsic defense mechanisms [50].

The reduction in placental O-GlcNAcylation extends beyond nutrient availability. Diverse studies have shown that protein O-GlcNAcylation regulates innate immune cell function, such as the NF-κB pathway [19,51,52]. Moreover, altered placental OGT expression can impact long-term metabolic and neurodevelopmental programming. The reduced OGT expression in a placental-specific OGT knockout mouse model altered the expression patterns of important hypothalamic genes involved in endocrine and anti-inflammatory signaling during development [25]. These alterations may mediate changes in placental endocrine function and differences in the transmission of important signals to fetal development. These are important mechanisms to be highlighted and understood, as children exposed *in utero* to ZIKV have greater risk of neurodevelopmental abnormalities in the first 18 months of life [53]. These findings reinforce the importance of monitoring the long-term neurological development of all newborns with exposure to ZIKV. While the placental HBP and O-GlcNAcylation are clearly altered in females, there were slight changes observed in males, again reinforcing the sex-specific characteristic of the response to ZIKV in this tissue. The specific sex difference observed in our work should be investigated since O-GlcNAcylation is identified as a potential mechanism related to sex-dependent neurodevelopment changes [26], and, as already mentioned, the relationship between infection by ZIKV and possible damage to several processes related to the nervous system is recurrent [37,54], but little explored in a sex-specific way.

In conclusion, our study highlights the importance of the placenta as a key organ in mediating gestational infection and points to relevant molecular alterations caused by maternal ZIKV infection. Collectively, our results suggest that female and male placentas adopt different strategies to cope with the altered metabolic state caused by ZIKV infection and that the alteration of placental nutrient transport and O-GlcNAcylation may be linked to the known effects of ZIKV infection. Our findings contribute to the understanding of alterations that could assist future preventive and therapeutic strategies and sex-specific approaches of individuals who had their mothers infected.

## Materials and methods

### Virus

The ZIKV strain PE243 (ZIKVPE243; GenBank ref. number KX197192) was kindly provided by Dr Ernesto T. Marques (University of Pittsburgh, PA). Viruses were propagated in C6/36 cells and titrated by plaque assays in Vero cells, as previously described (Coelho, et al., 2017). Conditioned medium from noninfected C6/36 cells cultured under the same conditions was used as a control.

### Experimental design

Female C57BL/6 mice aged between 8 and 10 weeks were mated with male mice, and on the day after mating, copulation was confirmed by visualization of the vaginal plug and considered gestational day 0.5 (GD0.5). Dam’s weight gain was monitored for confirmation of pregnancy. At GD 12.5, dams with a weight gain greater than 3 g were considered pregnant and randomly distributed into the following experimental groups: control (received an intravenous injection (i.v.) of supernatant from noninfected C6/36 cells, n=15), Low Dose (received an i.v. with 10^3^ plaque-forming units (PFU) of ZIKVPE243, n=15), and High Dose (received an i.v. with 5×10^7^ PFU of ZIKVPE243, n=15) ZIKV. The i.v. injections were performed at GD12.5, and virus preparation was undertaken as previously described [7,55]. At 18.5, all animals were weighed and euthanized with 5% isoflurane by inhalation. All placental/fetal units were dissected, collected and weighed. Fetal tails were kept for sex determination by detection of the *Sx* gene using specific primers (Forward: 5’- GATGATTTGAGTGGAAATGTGAGGTA-3’; Reverse: 3’- GAATACAAATATCCGTACGTGGTACAT-5’) [56].

Placental specimens were stored in liquid nitrogen for further qPCR and western blotting analyses. For the MALDI-IMS analysis, samples were frozen in acetone cooled with dry ice and subsequently kept in a -80 freezer. For morphology and protein expression/localization analysis, the tissues were fixed overnight in buffered paraformaldehyde (4%, Sigma‒Aldrich, Brazil). The experimental design is illustrated in Fig 1A. Animals were kept in a controlled temperature room (23°C) with a light/dark cycle of 12 hours and had access to water and food *ad libitum*. All procedures were approved by the Animal Care Committee (CEUA-036/16, 128/22, A7/20-036-16) of the Federal University of Rio de Janeiro.

### Histological and immunohistochemistry analysis

Previously fixed placental fragments were subjected to dehydration (increasing ethanol concentrations; ISOFAR, Brazil) and diaphanization with xylol (ISOFAR, Brazil). The fragments were embedded in paraffin (Histopar, Easypath, Brazil) and sectioned (5 µm thickness) using a Rotatory Microtome CUT 5062 (Slee Medical GmbH, Germany). Sections were submitted to the following techniques and performed as previously described [57]:

#### Periodic acid-Schiff (PAS) staining

Histological sections were submitted to diaphanization with three xylol baths and hydrated with three decreasing concentrations of ethanol. The sections were oxidized for 15 min. with 0.5% periodic acid (Sigma‒Aldrich, USA), washed in flowing water and incubated for 10 min. with Schiff’s reagent (Merck Millipore, Germany) at room temperature.

#### Immunochemistry of nutrient transporters in the placenta

The sections were submitted to diaphanization with three xylol baths and hydrated with three decreasing concentrations of ethanol. Subsequently, sections were exposed for 30 min to 3% hydrogen peroxide, followed by microwave antigenic recovery in Tris-EDTA (pH=9) and sodium citrate (pH=6) buffers (15 min. for Tris-EDTA buffer and 8 min. for citrate buffer). Sections were washed in PBS+ 0.2% Tween and incubated in 3% BSA in PBS for 1 hour to block nonspecific binding sites. Sections were then incubated with the primary antibody (Supplementary Table 1) overnight at 4°C. On the day after, sections were incubated with the biotin-conjugated secondary antibody (SPD-060, Spring Bioscience, USA) for 1 h at room temperature. Slides were incubated with streptavidin (SPD-060 – Spring Bioscience, USA) for 1 h and with 3,3-diaminobenzidine (DAB) (SPD-060 – Spring Bioscience, USA).

After PAS and immunochemistry procedures, all sections were counterstained with hematoxylin, followed by dehydration in alcohol and xylol. The sections were then mounted with coverslips using Entellan (Merck, Germany). Digital images were obtained using a high-resolution Olympus DP72 (Olympus Corporation, Japan) camera coupled to an Olympus BX53 light microscope (Olympus Corporation, Japan). The PAS-stained sections were used to analyze the area of each placenta of the rodent placental zones, labyrinth (Lz) and junctional (Jz) zones by free-drawing using ImageJ software (National Institutes of Health, USA). For immunochemistry quantification of Glut1 and Snat2, fifteen digital images (40X) from different fields were captured in each placenta zone and analyzed with Image-Pro Plus, version 5.0 software (Media Cybernetics, USA) mask tool. At each staining, a total of 150 digital images were analyzed per group (75 Lz images and 75 Jz images). Hence, 450 images were analyzed by target protein including all experimental groups. The percentage of viable tissue area was considered upon the exclusion of negative spaces. All negative controls were performed with the omission of the primary antibody. Analysis were performed as previously described [7]. All analyses were conducted blind to the group.

### RT‒qPCR

Placental RNA was extracted using TRIzol following the manufacturer’s protocol (TRIzol Reagent; Life Technologies, USA). The concentration of RNA was assessed using a NanoPhotometer (Implen, Munchen, Germany). The quantification and RNA purity were evaluated by reading in a nanophotometer, considering the relations A260/280 ∼2.0 and A260/230 ∼2.0-2.2. (Montreal Biotech Inc., Canada). A total of 1 μg per sample was reverse transcribed using the High-Capacity cDNA Reverse Transcription Kit (Applied Biosystems, USA) according to the manufacturer’s instructions. qPCR was performed to evaluate mRNA expression quantification using gene-specific primer pairs (Supplementary Table 2) and EVAGREEN (Solis Byodine, USA). The standard thermal cycling protocol was conducted as follows: combined initial denaturation at 50 °C (2 min) and 95° C (10 min), followed by 40 cycles of denaturation at 95 °C (15 s), annealing at 60 °C (30 s) and extension at 72 °C (45 s), using QIAquant 96 2plex (QIAGEN, Germany). Relative expression was calculated using the 2-ΔΔCt method [58], and genes of interest were normalized to the geometric mean expression of 3 reference genes (*Ywhaz, PPib, ß-actin*).

### Western blotting

Placental protein extraction was performed on frozen tissue homogenized using commercial RIPA lysis buffer (Thermo Scientific, US) supplemented with phosphatase inhibitors (5 mM Na4P2O7, 50 mM NaF, 5 mM Na3VO4), protease inhibitor mixture (Sigma‒ Aldrich), and OGA inhibitor (PugNAc, 1 M). The protein concentration was determined using the bicinchoninic acid protein assay (Thermo Fisher Scientific, EUA). Lysates were separated by sodium dodecyl sulfate‒polyacrylamide gel electrophoresis (SDS‒PAGE) and transferred onto nitrocellulose membranes (Bio-Rad Laboratories, US) using a semidry technique (semidry blotter, Invitrogen). The membrane washing steps were performed with Tris-buffered saline tween (TBST) after the following blocking and incubation steps. Blocking was performed with 3% BSA in phosphate-buffered saline (1x) with 0.1% Tween (1X Tris-buffered saline, 0.1% Tween - TBS-T) for 1 h at room temperature. The membrane was incubated with primary antibody (Supplementary Table 1) in 3% BSA + TBS-T overnight. The next day, the membranes were washed and incubated with rabbit or mouse secondary antibodies diluted in TBST for 60 minutes. ß-actin was used as a loading control. Protein bands were visualized using Scientific SuperSignal West Femto enhanced chemiluminescence (ECL) substrate (Thermo Scientific, US) and captured with ImageQuant LAS 4000.

### MALDI-IMS

The analysis of specific biomolecules enriched in murine placentas exposed to ZIKV was performed using matrix-assisted laser desorption/ionization imaging mass spectrometry (MALDI-IMS). We used the mass/charge ratio (m/z) to analyze our biomolecules of interest: glucose-6-phosphate (m/z= 259.0232), glutamine (m/z= 145.0618) and UDP-GlCNAc (m/z= 606.3824).

The presence of the biomolecules was analyzed in 5 placentas (randomly picked) per experimental group. Cryosectioning of previously frozen placentas was performed with 10-mm-thick sections using a cryostat (Leica CM1860-UV, Leica Biosystems, Nussloch, Germany). Then, the slices were added to glass slides coated with ITO (indium tin oxide) and conditioned at -80°C until the time of analysis. The slides were removed from -80°C and allowed to reach room temperature in the desiccator until the time of deposition of the matrix. Subsequently, deposition of the matrix was carried out. The matrix solution used was 10 mg/mL 9-aminoacridine (Sigma) in 80% methanol and 20% water. The application was performed with spray with an application flow of 800 μL/h and N2 pressure of 12.5 psi for 20 minutes. The analysis was performed with a Solarix XR mass spectrometer (FT-ICR, Bruker) using negative polarity, mass range from 75 to 1200 m/z, laser focus medium mode and power make laser at 35%. Two hundred shots were fired per field. The data obtained in the spectrometry were normalized into TIC (total ion count).

### Statistics

Statistical analyses were performed using GraphPad Prism version 8 (GraphPad, La Jolla, CA, USA). A D’Agostino & Pearson normality test was used to evaluate normal distribution. Data for each sex were analyzed using one-way ANOVA followed by Tukey’s test. Outliers were identified using a Grubbs test. Differences were considered significant when p<0.05. Litter parameters were evaluated using the mean value of all fetuses and/or placentae in the litter per dam and not the individual conceptus [59,60]. The number of samples per group for each analysis is shown in each figure and described in the legends of figures and footnotes of tables.

## Acknowledgments

We would like to thank Ernesto T. Marques Jr. (Centro de Pesquisa Aggeu Magalhães, FIOCRUZ, PE) for providing ZIKV to the Institute of Microbiology Paulo de Góes.

---

**Supplementary figure 1.** Nutrient transport in the junctional zone of control and ZIKV-infected placentas. **(A)** Quantification and representative photomicrographs of Glut1 expression in the placental junctional zone by immunohistochemistry in female placentas (n=5). **(B)** Quantification and representative photomicrographs of Snat2 expression in the placental junctional zone by immunohistochemistry in female placentas (n=5). (**C)** Quantification and representative photomicrographs of Glut1 expression in the placental junctional zone by immunohistochemistry in male placentas (n=5). **(D)** Quantification and representative photomicrographs of Snat2 expression in the placental junctional zone by immunohistochemistry in male placentas (n=5). Data are displayed as the mean + SEM. and analyzed by one-way ANOVA with Tukey post hoc comparisons.

**Supplementary table 1.** List of primary antibodies used in this study.

| Primary antibody | Host | Manufacturer, catalog number | Dilution |
| --- | --- | --- | --- |
| Actin | *Mouse* | Sigma–Aldrich, A5441 | 1/2000 |
| Glut1 | *Rabbit* | Invitrogen, PA5-16793 | 1/2000 |
| Snat2 | *Rabbit* | Bioss, bs-12125R | 1/100 |
| Gfat1 | *Rabbit* | Santa Cruz Biotechnology, SC- 134894 | 1/1000 |
| Gfat2 | *Rabbit* | Santa Cruz Biotechnology, SC- 134710 | 1/1000 |
| OGT | *Mouse* | Santa Cruz Biotechnology, (F-12), SC-74546 | 1/2000 |
| OGA | *Rabbit* | Santa Cruz Biotechnology, 345 | 1/2000 |
| O-GlcNAc | *Mouse* | Santa Cruz Biotechnology, RL-2, SC59624 | 1/1000 |

**Supplementary table 2.**
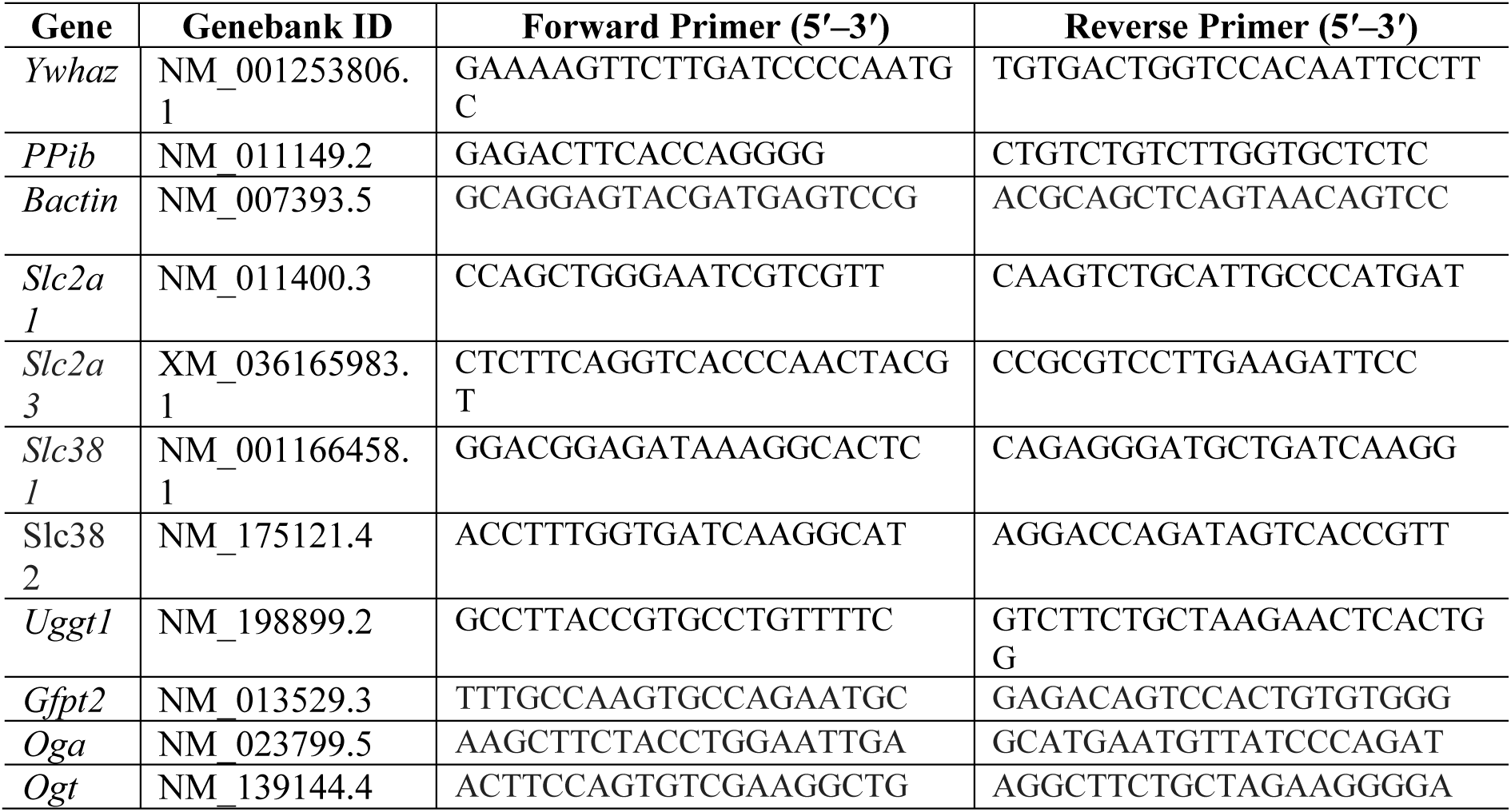
List of primers used for quantitative RT‒PCR

## References

1. Oliveira Melo AS, Malinger G, Ximenes R, Szejnfeld PO, Alves Sampaio S, et al. (2016) Zika virus intrauterine infection causes fetal brain abnormality and microcephaly: tip of the iceberg? Ultrasound Obstet Gynecol 47: 6–7.

2. Messina JP, Kraemer MU, Brady OJ, Pigott DM, Shearer FM, et al. (2016) Mapping global environmental suitability for Zika virus. Elife 5.

3. Chen Q, Gouilly J, Ferrat YJ, Espino A, Glaziou Q, et al. (2020) Metabolic reprogramming by Zika virus provokes inflammation in human placenta. Nat Commun 11: 2967.

4. Thaker SK, Chapa T, Garcia G, Jr., Gong D, Schmid EW, et al. (2019) Differential Metabolic Reprogramming by Zika Virus Promotes Cell Death in Human versus Mosquito Cells. Cell Metab 29: 1206–1216 e1204.

5. Yau C, Low JZH, Gan ES, Kwek SS, Cui L, et al. (2021) Dysregulated metabolism underpins Zika-virus-infection-associated impairment in fetal development. Cell Rep 37: 110118.

6. Creisher PS, Lei J, Sherer ML, Dziedzic A, Jedlicka AE, et al. (2022) Downregulation of transcriptional activity, increased inflammation, and damage in the placenta following in utero Zika virus infection is associated with adverse pregnancy outcomes. Front Virol 2.

7. Andrade CBV, Monteiro VRS, Coelho SVA, Gomes HR, Sousa RPC, et al. (2021) ZIKV Disrupts Placental Ultrastructure and Drug Transporter Expression in Mice. Front Immunol 12: 680246.

8. Winterhager E, Gellhaus A (2017) Transplacental Nutrient Transport Mechanisms of Intrauterine Growth Restriction in Rodent Models and Humans. Front Physiol 8: 951.

9. Coan PM, Vaughan OR, Sekita Y, Finn SL, Burton GJ, et al. (2010) Adaptations in placental phenotype support fetal growth during undernutrition of pregnant mice. J Physiol 588: 527–538.

10. Higgins JS, Vaughan OR, Fernandez de Liger E, Fowden AL, Sferruzzi-Perri AN (2016) Placental phenotype and resource allocation to fetal growth are modified by the timing and degree of hypoxia during mouse pregnancy. J Physiol 594: 1341–1356.

11. Lager S, Powell TL (2012) Regulation of nutrient transport across the placenta. J Pregnancy 2012: 179827.

12. Vaughan OR, Maksym K, Silva E, Barentsen K, Anthony RV, et al. (2021) Placenta-specific Slc38a2/SNAT2 knockdown causes fetal growth restriction in mice. Clin Sci (Lond) 135: 2049–2066.

13. Gaccioli F, Lager S (2016) Placental Nutrient Transport and Intrauterine Growth Restriction. Front Physiol 7: 40.

14. Li P, He L, Lan Y, Fang J, Fan Z, et al. (2022) Intrauterine Growth Restriction Induces Adulthood Chronic Metabolic Disorder in Cardiac and Skeletal Muscles. Front Nutr 9: 929943.

15. Chandrasiri UP, Chua CL, Umbers AJ, Chaluluka E, Glazier JD, et al. (2014) Insight into the pathogenesis of fetal growth restriction in placental malaria: decreased placental glucose transporter isoform 1 expression. J Infect Dis 209: 1663–1667.

16. Vota D, Torti M, Paparini D, Giovannoni F, Merech F, et al. (2021) Zika virus infection of first trimester trophoblast cells affects cell migration, metabolism and immune homeostasis control. J Cell Physiol 236: 4913–4925.

17. Hart GW, Slawson C, Ramirez-Correa G, Lagerlof O (2011) Cross talk between O-GlcNAcylation and phosphorylation: roles in signaling, transcription, and chronic disease. Annu Rev Biochem 80: 825–858.

18. Hart B, Morgan E, Alejandro EU (2019) Nutrient sensor signaling pathways and cellular stress in fetal growth restriction. J Mol Endocrinol 62: R155–R165.

19. Quik M, Hokke CH, Everts B (2020) The role of O-GlcNAcylation in immunity against infections. Immunology 161: 175–185.

20. Shi Y, Yan S, Shao GC, Wang J, Jian YP, et al. (2022) O-GlcNAcylation stabilizes the autophagy-initiating kinase ULK1 by inhibiting chaperone-mediated autophagy upon HPV infection. J Biol Chem 298: 102341.

21. Zeng Q, Zhao RX, Chen J, Li Y, Li XD, et al. (2016) O-linked GlcNAcylation elevated by HPV E6 mediates viral oncogenesis. Proc Natl Acad Sci U S A 113: 9333–9338.

22. Wang X, Lin Y, Liu S, Zhu Y, Lu K, et al. (2020) O-GlcNAcylation modulates HBV replication through regulating cellular autophagy at multiple levels. FASEB J 34: 14473–14489.

23. Wang Q, Fang P, He R, Li M, Yu H, et al. (2020) O-GlcNAc transferase promotes influenza A virus-induced cytokine storm by targeting interferon regulatory factor-5. Sci Adv 6: eaaz7086.

24. Angelova M, Ortiz-Meoz RF, Walker S, Knipe DM (2015) Inhibition of O-Linked N-Acetylglucosamine Transferase Reduces Replication of Herpes Simplex Virus and Human Cytomegalovirus. J Virol 89: 8474–8483.

25. Howerton CL, Bale TL (2014) Targeted placental deletion of OGT recapitulates the prenatal stress phenotype including hypothalamic mitochondrial dysfunction. Proc Natl Acad Sci U S A 111: 9639–9644.

26. Howerton CL, Morgan CP, Fischer DB, Bale TL (2013) O-GlcNAc transferase (OGT) as a placental biomarker of maternal stress and reprogramming of CNS gene transcription in development. Proc Natl Acad Sci U S A 110: 5169–5174.

27. Moore M, Avula N, Jo S, Beetch M, Alejandro EU (2021) Disruption of O-Linked N-Acetylglucosamine Signaling in Placenta Induces Insulin Sensitivity in Female Offspring. Int J Mol Sci 22.

28. Nugent BM, O’Donnell CM, Epperson CN, Bale TL (2018) Placental H3K27me3 establishes female resilience to prenatal insults. Nat Commun 9: 2555.

29. Yang YR, Jang HJ, Lee YH, Kim IS, Lee H, et al. (2015) O-GlcNAc cycling enzymes control vascular development of the placenta by modulating the levels of HIF-1alpha. Placenta 36: 1063–1068.

30. Nem de Oliveira Souza I, Frost PS, Franca JV, Nascimento-Viana JB, Neris RLS, et al. (2018) Acute and chronic neurological consequences of early-life Zika virus infection in mice. Sci Transl Med 10.

31. Paul AM, Acharya D, Neupane B, Thompson EA, Gonzalez-Fernandez G, et al. (2018) Congenital Zika Virus Infection in Immunocompetent Mice Causes Postnatal Growth Impediment and Neurobehavioral Deficits. Front Microbiol 9: 2028.

32. Szaba FM, Tighe M, Kummer LW, Lanzer KG, Ward JM, et al. (2018) Zika virus infection in immunocompetent pregnant mice causes fetal damage and placental pathology in the absence of fetal infection. PLoS Pathog 14: e1006994.

33. Suzukawa AA, Zanluca C, Jorge NAN, de Noronha L, Koishi AC, et al. (2020) Downregulation of IGF2 expression in third trimester placental tissues from Zika virus infected women in Brazil. J Infect 81: 766–775.

34. Nardozza LM, Caetano AC, Zamarian AC, Mazzola JB, Silva CP, et al. (2017) Fetal growth restriction: current knowledge. Arch Gynecol Obstet 295: 1061–1077.

35. Burton GJ, Jauniaux E (2018) Pathophysiology of placental-derived fetal growth restriction. Am J Obstet Gynecol 218: S745–S761.

36. Sferruzzi-Perri AN, Camm EJ (2016) The Programming Power of the Placenta. Front Physiol 7: 33.

37. Andrade TA, Fahel JS, de Souza JM, Terra AC, Souza DG, et al. (2022) In Utero Exposure to Zika Virus Results in sex-Specific Memory Deficits and Neurological Alterations in Adult Mice. ASN Neuro 14: 17590914221121257.

38. Sherer ML, Lemanski EA, Patel RT, Wheeler SR, Parcells MS, et al. (2021) A Rat Model of Prenatal Zika Virus Infection and Associated Long-Term Outcomes. Viruses 13.

39. Salazar-Petres E, Pereira-Carvalho D, Lopez-Tello J, Sferruzzi-Perri AN (2022) Placental structure, function, and mitochondrial phenotype relate to fetal size in each fetal sex in micedagger. Biol Reprod 106: 1292–1311.

40. Monteiro VRS, Andrade CBV, Gomes HR, Reginatto MW, Imperio GE, et al. (2022) Mid-pregnancy poly(I:C) viral mimic disrupts placental ABC transporter expression and leads to long-term offspring motor and cognitive dysfunction. Sci Rep 12: 10262.

41. Sakata M, Kurachi H, Imai T, Tadokoro C, Yamaguchi M, et al. (1995) Increase in human placental glucose transporter-1 during pregnancy. Eur J Endocrinol 132: 206–212.

42. Das UG, Sadiq HF, Soares MJ, Hay WW, Jr., Devaskar SU (1998) Time-dependent physiological regulation of rodent and ovine placental glucose transporter (GLUT-1) protein. Am J Physiol 274: R339–347.

43. Luscher BP, Marini C, Joerger-Messerli MS, Huang X, Hediger MA, et al. (2017) Placental glucose transporter (GLUT)-1 is down-regulated in preeclampsia. Placenta 55: 94–99.

44. Yao G, Zhang Y, Wang D, Yang R, Sang H, et al. (2017) GDM-Induced Macrosomia Is Reversed by Cav-1 via AMPK-Mediated Fatty Acid Transport and GLUT1-Mediated Glucose Transport in Placenta. PLoS One 12: e0170490.

45. Borges MH, Pullockaran J, Catalano PM, Baumann MU, Zamudio S, et al. (2019) Human placental GLUT1 glucose transporter expression and the fetal insulin-like growth factor axis in pregnancies complicated by diabetes. Biochim Biophys Acta Mol Basis Dis 1865: 2411–2419.

46. Yang M, Li H, Rong M, Zhang H, Hou L, et al. (2022) Dysregulated GLUT1 may be involved in the pathogenesis of preeclampsia by impairing decidualization. Mol Cell Endocrinol 540: 111509.

47. Langdown ML, Sugden MC (2001) Enhanced placental GLUT1 and GLUT3 expression in dexamethasone-induced fetal growth retardation. Mol Cell Endocrinol 185: 109–117.

48. Sibiak R, Ozegowska K, Wender-Ozegowska E, Gutaj P, Mozdziak P, et al. (2022) Fetomaternal Expression of Glucose Transporters (GLUTs)-Biochemical, Cellular and Clinical Aspects. Nutrients 14.

49. Chiaradonna F, Ricciardiello F, Palorini R (2018) The Nutrient-Sensing Hexosamine Biosynthetic Pathway as the Hub of Cancer Metabolic Rewiring. Cells 7.

50. Zhang S, Regnault TR, Barker PL, Botting KJ, McMillen IC, et al. (2015) Placental adaptations in growth restriction. Nutrients 7: 360–389.

51. Dong H, Liu Z, Wen H (2022) Protein O-GlcNAcylation Regulates Innate Immune Cell Function. Front Immunol 13: 805018.

52. Dela Justina V, Goncalves JS, de Freitas RA, Fonseca AD, Volpato GT, et al. (2017) Increased O-Linked N-Acetylglucosamine Modification of NF-KappaB and Augmented Cytokine Production in the Placentas from Hyperglycemic Rats. Inflammation 40: 1773–1781.

53. Mulkey SB, Arroyave-Wessel M, Peyton C, Bulas DI, Fourzali Y, et al. (2020) Neurodevelopmental Abnormalities in Children With In Utero Zika Virus Exposure Without Congenital Zika Syndrome. JAMA Pediatr 174: 269–276.

54. Trus I, Udenze D, Cox B, Berube N, Nordquist RE, et al. (2019) Subclinical in utero Zika virus infection is associated with interferon alpha sequelae and sex-specific molecular brain pathology in asymptomatic porcine offspring. PLoS Pathog 15: e1008038.

55. Coelho SVA, Neris RLS, Papa MP, Schnellrath LC, Meuren LM, et al. (2017) Development of standard methods for Zika virus propagation, titration, and purification. J Virol Methods 246: 65–74.

56. Davalieva K, Dimcev P, Efremov GD, Plaseska-Karanfilska D (2006) Non-invasive fetal sex determination using real-time PCR. J Matern Fetal Neonatal Med 19: 337–342.

57. Fontes KN, Reginatto MW, Silva NL, Andrade CBV, Bloise FF, et al. (2019) Dysregulation of placental ABC transporters in a murine model of malaria-induced preterm labor. Sci Rep 9: 11488.

58. Livak KJ, Schmittgen TD (2001) Analysis of relative gene expression data using real-time quantitative PCR and the 2(-Delta Delta C(T)) Method. Methods 25: 402–408.

59. Connor KL, Kibschull M, Matysiak-Zablocki E, Nguyen TTN, Matthews SG, et al. (2020) Maternal malnutrition impacts placental morphology and transporter expression: an origin for poor offspring growth. J Nutr Biochem 78: 108329.

60. Bloise E, Lin W, Liu X, Simbulan R, Kolahi KS, et al. (2012) Impaired placental nutrient transport in mice generated by in vitro fertilization. Endocrinology 153: 3457–3467.

