## Supplementary figures and images for "Sex-specific effect of antenatal Zika virus infection on murine fetal growth, placental nutrient transporters, and nutrient sensor signaling pathways"

### Supplementary figure 1

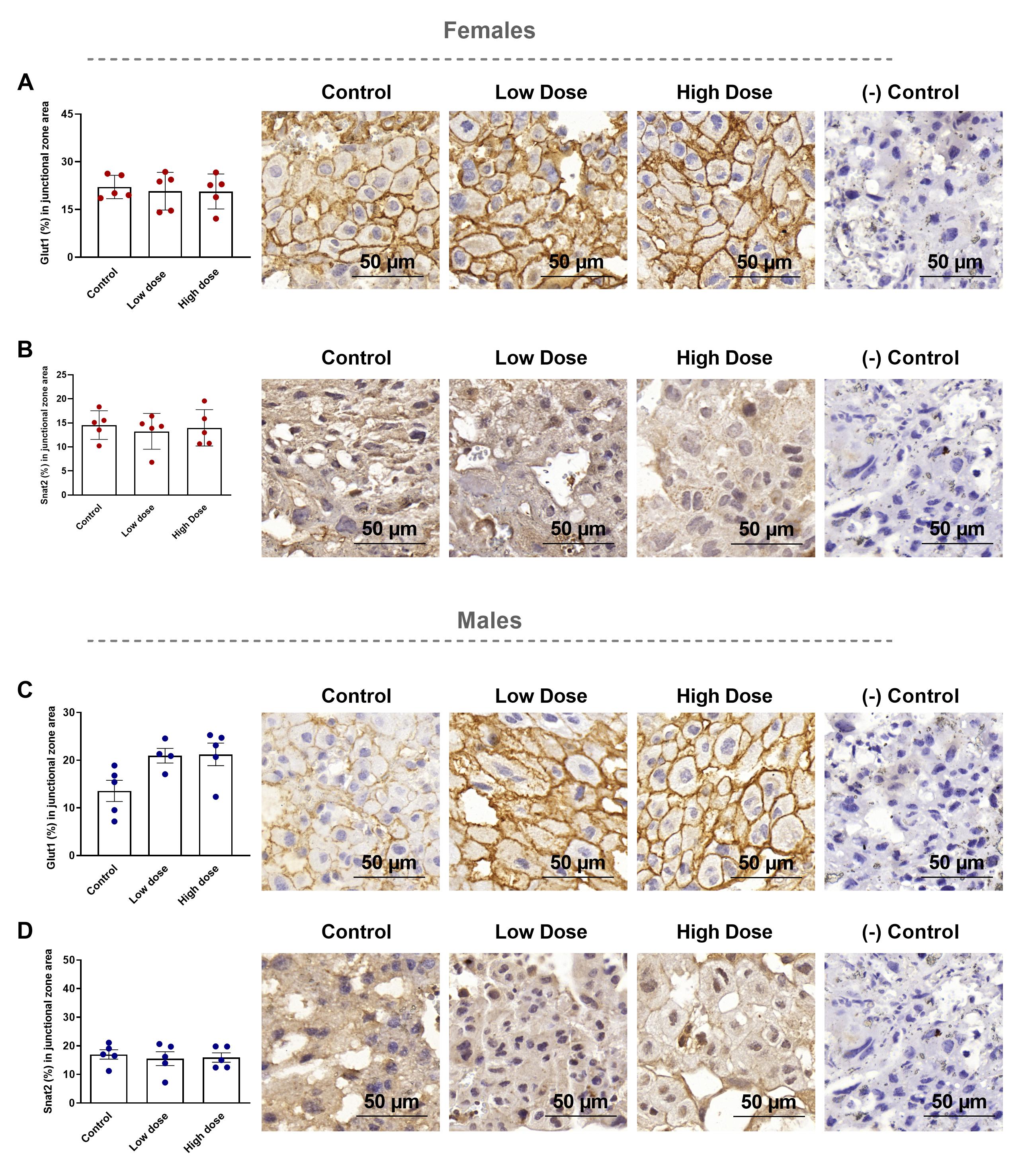
